# Fast and Scalable Genome-Wide Inference of Local Tree Topologies from Large Number of Haplotypes Based on Tree Consistent 𝒫ℬ𝒲𝒯 Data Structure

**DOI:** 10.1101/542035

**Authors:** Vladimir Shchur, Liliia Ziganurova, Richard Durbin

## Abstract

Estimation of the relationship between DNA sequences is one of the most important problems in genomics. Understanding these relationships is central to demographic inference, correction of population structure in GWAS, identifying signals of selection etc. The data structure containing the full information about sample genealogy is called the ancestral recombination graph 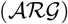. However, 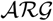 inference is a very difficult problem, not least due to a very complex state space. In this work we describe a new approach for fast and scalable generation of local tree topologies relating large numbers of haplotypes. Our method is closely related to the estimation of 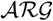, and captures both local and global properties of an 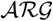. It is based on a data structure which we call tree consistent 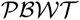, a modification of 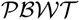 data structure introduced by R. Durbin (2014). We also explore some methods to estimate the quality of the generated tree topologies and to make inferences based on them. At the end we discuss a probabilistic model which could potentially lead to the estimation of 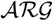 node times.

## 1. Introduction

During recent years the size of sequencing DNA data increases extensively. This continual data growth in principal enables deeper understanding of population structure and genetic nature of biological traits. We need new fast algorithms and tools to make possible data processing and inference. R. Durbin [2] suggested a new way to represent a set of genome sequences or haplotypes called the “positional Burrows-Wheeler transform” or shortly 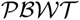. This representation allows very fast and effective data compression and haplotype matching, and underlies the rapid genotype imputation process used by the Sanger Institute Haplotype Reference Consortium imputation server [8]. The important feature of the 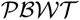-based algorithms is their scalability: they are linear or close to linear in the amount of data.

The complete information about sample history is contained in the Ancestral Recombination Graph, shortly 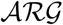 [4]. Such a graph represents a genealogical network which shows in detail a genealogy for every locus, as well as recombinations that link these genealogies. Suppose that there is a set of prealligned genomic sequences. At every site all these sequences are genealogically related by a *local tree*. The corresponding 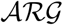 can be represented as a set of local trees. For completeness, the recombinations which transform one tree to another also should be specified. In fact they appear as *prune-and-regraft* operations on local trees [9]: an edge of a local tree is pruned, resulting in a subtree which is cut from the rest of the tree, and then this subtree is regrafted somewhere on the remaining tree. The regrafting point must be higher (earlier in time) than the pruning point.

In this paper we present an alternative form of the 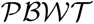, which we call the “tree consistent 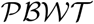” or shortly 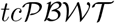. It follows the initial philosophy of 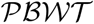 and remains linear, though it uses additional information about which site at an allele is ancestral. The data compression rate of 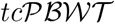 slightly improves compared to the original 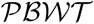 which means that 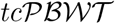 has more internal structure than 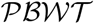. This implies that 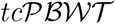 captures genomic structure better than 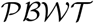. We will establish the relation between the 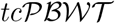 structure of the haplotype set and the subtrees in the local coalescent tree that relate the haplotypes at each position in the genome. Based on this we will be able to identify recombination events and hence build a representation of an ancestral recombination graph (ARG) which relates the full set of haplotypes to each other. Alternative choices in selecting recombination events will lead to alternative downstream ARGs. Potentially this ambiguity allows us to sample genealogical histories for use in statistical inference.

As was already mentioned, the 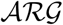 contains all the genealogical information of a set of haplotypes, which makes the 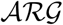 a very valuable object. However the inference of the 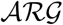 is a very challenging problem because it is underdetermined and the corresponding state space is enormous. To simplify the problem we start with the topology construction while ignoring edge lengths. We will show that the 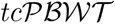 algorithm will find the correct topology of the tree in case of the perfect phylogeny (without recombinations, and with at most one mutation at each site).

We also establish a probabilistic model for the 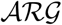 topology. We will state the problem of time estimation for 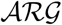 nodes for a given topology as a problem of mathematical programming. Currently we did not succeed in applying this probabilistic model to the 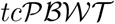 data structure.

To our knowledge, the most scalable existing methods to reconstruct possible 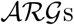 are *MARGarita* by Minichiello and Durbin (2006) [10] and *ARGweaver* by Rasmussen et al. (2014) [11]. *MARGarita* is an heuristic algorithm; whereas *ARGweaver* infers 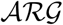 under the Sequential Markov Coalescent (SMC), which is an approximation introduced by McVean and Cardin (2005) [9] of the coalescent with recombination process [5] [3]. SMC models 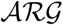 as a process with Markovian structure with similar properties compared to the coalescence with recombination. *ARGweaver* can be applied to dozens or even a few hundred genomes, it is linear in the number of sites, quadratic in the number of haplotypes and quadratic in the number of discretised time points. Its main idea is to infer the 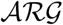 for the set of haplotypes {*h*_0_, *h*_1_,…, *h_i_*} by adding the haplotype *h_i_* to the preinfered and then fixed 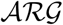 for {*h*_0_, *h*_1_, …, *h*_*i*−1_}.

Another possible approach used in [13] is to infer only topologies of local trees. Basically, one might want to find for each region the longest fully compatible regions and to use this information to build a tree. But trees in adjacent regions share a lot of structure, which is used for further improvement of the inference of trees. Our approach uses a similar idea, and our 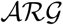 data structure insures that the inference is fast and scalable.

In this paper we suggest an approach for fast generation of 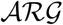 topologies from genomic data. We also suggest data structures which allow to achieve even faster performance of the method. This data structures can be also used for efficient and as a compact data storing of genomic variation and genealogical histories of a sample. We will demonstrate the scalability of our method. We will also discuss the consistency of our approach with the true underlying 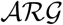 by simulations. Finally, we present some exploratory analysis of 1000 Genomes project Phase 3 data using the 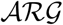 built from chromosome 20.

Further we discuss a probabilistic model for 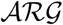 topologies. We made two attempts to apply this model to inferred topologies to estimate coalescent times and accuracy of local trees, but we did not get reliable results. Though, if we were able to find a solution, it would allow to generate samples of 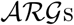 compatible with the data. Hence potentially it would enable the possibility to integrate over a sample of 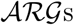 in order to use it for an inference of different features and statistics (both topological and related to coalescent times)

## 2. Notations and basic concepts

### 2.1. Input data

Firstly, let us give definitions and formalisation of the main objects in the paper. *A haplotype* is a DNA sequence ordered along the genome. This is a sequence of values at genetic sites that vary within a sample of individuals due to a mutation since their common ancestor. Formally, a haplotype of length *N* is a binary string *h* ∈ {0, 1}^*N*^. We denote a set of prealigned haplotypes by *H* = {*h*_0_, *h*_1_, …, *h*_*M*−1_}. The binary symbols obtain the following sense: 0 stands for ancestral allele (the value present in the common ancestor of the samples), 1 is for derived allele created by a mutation. A value of *h* at a particular site *k* is referred by *h*[*k*]. For this paper we will work in the infinite sites model in which there is only mutation at each site. Some algorithms we give can be extended to the case where there may be multiple mutations at a site but that is future work.

A set of haplotypes *H* can be considered as a matrix where any haplotype is a row, or a string, and a site *H_k_* = {*h*_0_[*k*], *h*_1_[*k*], …, *h*_*M*−1_[*k*]} is a column.

### 2.2. Ancestral Recombination Graph

The coalescent with recombination is a probabilistic model describing the distribution of genealogies of a sample of haplotypes [5] [3]. As for the standard coalescent, the ancestral lineages start at the present and go backwards in time. There are two types of events that may happen with a lineage: either two lineages coalesce (coalescent node) into one lineage or a lineage is split by recombination (recombination node) into two lineages. The 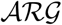 is a graph representing this process. As a data structure, 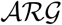 is a directed acyclic graph with a time function (consisting with the orientation of edges) and with additional information at recombination nodes on breakpoints. For every site an 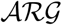 embeds a genealogical local tree.

We remind that a tree is a graph without cycles. A rooted tree is an oriented tree where one vertex (*the root*) has in-degree 0 and all other vertices have in-degree 1. The vertices with out-degree 0 are called *leaves*. The vertices which are not leaves are *internal nodes*.

A genealogical local tree is a rooted tree. A time is assigned to every vertex of this tree. Usually we consider time in the backward direction so that the time assigned to the leaves is 0 and the time of a parent vertex is always larger than the time of the child vertex.

Subtree prune-and-regraft operation, or SPR in short, is a transformation of a tree which consists of following steps:

- cutting (pruning) an edgeg of the tree at a certain point;
- if the parental node of the edge was binary before pruning, it is removed;
- the pruned subtree is regrafted at any place on the tree above the cut point (equivalently, earlier in the past).

*Perfect phylogeny* means that there is at most one mutation at every site of genome. We call a site *H* compatible with a tree *T* if it is possible to add at most one mutation on this tree to explain *H*. In other words, there is a subtree *T^H^* of the tree *T* such that all haplotypes in *T_H_* carry the derived allele at the site and all haplotypes in the complementary *T* \ *T_H_* have the ancestral allele.

### 2.3. 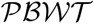 data structure

We briefly present the 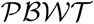 representation [2] of a set *H* of haplotypes. Basically, 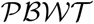 moves along the genome and reorders *H* site by site following some particular rules. Suppose that *H* is already ordered by all sites before *k* and the order is determined by the permutation *a*_*k*−1_. Now consider a permutation of *a*_*k*−1_(*H_k_*) (this is a permuted site *k* of *H*) which sends all 1s to the bottom of column *k*, while keeping the relative order between haplotypes sharing the same value at the site. More formally, if *h*_*a*_*k*−1_(*i*)_[*k*] is 0, then we set *a_k_*(*i*) to be the number of haplotypes *h_j_* such that *h_j_*[*k*] = 0 and *a*_*k*−1_(*j*) < *a*_*k*−1_(*i*). Similarly, if *h*_*a*_*k*−1_(*i*)_[*k*] is 1, then we set *a_k_*(*i*) to be the number of haplotypes *h_j_* such that *h_j_*[*k*] = 1 and *a*_*k*−1_(*j*) < *a*_*k*−1_(*i*) plus the total number of haplotypes for which *h_j_* [*k*] = 0.

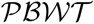 is a very fast operation. At a given site, this procedure corresponds to reordering of H into reverse lexicographical prefix order. This means that one considers prefixes of haplotypes relatively to a given site. The prefixes are read from right to left, which explains why we refer to the *reverse* order, and are ordered as the words in a dictionary, lexicographically. This property guarantees that the adjacent haplotypes share the local set-maximal left-match: consider again prefixes at the site, select one of them and compare to the rest. Then the prefix which shares the longest match at its end with the chosen one, in 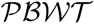 appears just after it. We denote the length of the shared left-match at site *k* between two haplotypes *h_i_* and *h_j_* by *d_k_*(*h_i_,h_j_*).

Generally, the longer is the local match between two haplotypes, the more locally related are they. So the adjacent haplotypes in 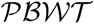 tend to have the same value at the next position. Indeed, in practice a permuted site *a*_*k*−1_(*H_k_*) demonstrates a lot of structure and usually consists of long stretches with constant values within them. For this reason we call *d_k_* “a similarity function”: the larger is the value of *d_k_*, the more relation haplotypes tend to show. An important property of this similarity function is that

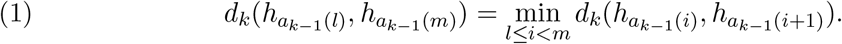

The similarity function allows 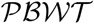 for example to be used for imputation by choosing similar haplotypes to impute from.

## 3. Compact tree and 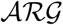 representation

In this section we will describe a fast and compact data structure which allows to achieve high efficiency of 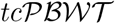 method. It also provides a compact representation of 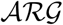, so it can be used as a data format for storing genealogy and genomic variation of a sample.

### 3.1. Planar ordering and compact tree representation

Firstly we formalise the intuitive approach of ordering leaves of a tree. Let us say that one wants to draw a genealogical tree of several individuals from the same generation on a sheet of paper. Two reasonable requirements for the drawing are to put all leaves of the tree on the same line and to represent it without self-intersections of edges. The corresponding order of the leaves on the line we call a *planar order* of a tree. This order is not unique and it is defined up to swapping subtrees rooted at the same internal node.

#### Definition 1.

*Planar order σ* of a tree *T* is an enumeration of its leaves which is defined in the following iterative process.

- **Initialization.** *S* is the set of all leaves, 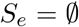 is an empty set. Choose the first leaf *h*_0_ randomly from *S*. Set *S_e_* = {*h*_0_}.
- **Recursion step.** Let *h_i_* be the last chosen (enumerated) leaf. Let *S_m_* be a subset of *S* \ *S_e_* of those leaves which minimise the distance to *h_i_* in the tree. Choose randomly *h*_*i*+1_ from *S_m_*. Set *S_e_* = *S_e_* ∪ {*h*_*i*+1_}.
- **Termination.** The process stops as soon as we enumerated all leaves of *T*.

A tree *T* is completely encoded by the planar order *σ*(0), *σ*(1), …, *σ*(*M* − 1) and distances between adjacent (in this order) leaves *d*(*h*_*σ*(0)_, *h*_*σ*(1)_), *d*(*h*_*σ*(1)_, *h*_*σ*(2)_), …, *d*(*h*_*σ*(*M*−2)_, *h*_*σ*(*M*−1)_) which we call the *distance vector*. For simplicity of notations denote *d*(*h*_*σ*(*i*−1)_, *h*_*σ*(*i*)_) by *d*_*σ*(*i*)_.

The distance between any two leaves *h*_*σ*(*j*)_ and *h*_*σ*(*k*)_ is given by the maximal value of the distance vector on the semi-open interval (*σ*(*j*),*σ*(*k*)] and can be computed by the following formula

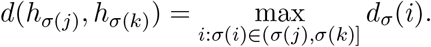

Now notice that referring to an internal node of a tree is equivalent to referring to a pair of leaves of a tree such that their most recent common ancestor (shortly, MRCA) is this node. Planar order provides such a correspondence between pairs of adjacent leaves and internal nodes. We encode an internal node *ν* by the value of the index *i* such that the node *ν* is *MRCA*(*h*_*σ*(*i*−1)_, *h*_*σ*(*i*)_). In case of a non-binary tree by convention we encode the node *ν* by the minimal value of *i* satisfying the previous condition (see Figure 1 for an example).

**Figure 1.**
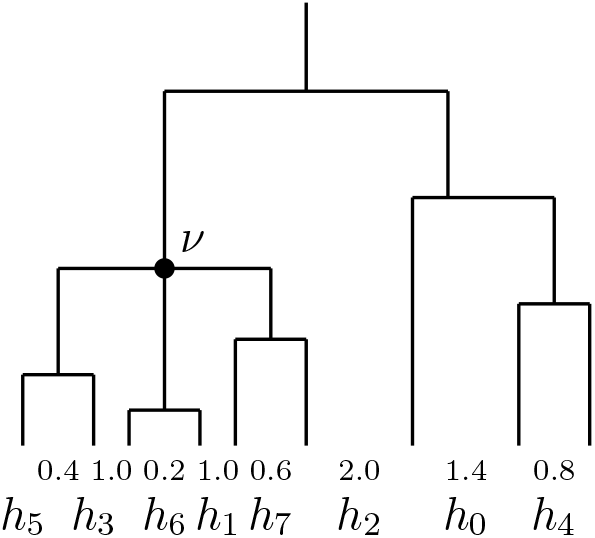
Planar order: the tree is encoded by vectors *σ* = (5, 3, 6, 1, 7, 2, 0, 4) and *d* = (0.4, 1.0, 0.2, 1.0, 0.6, 2.0,1.4, 0.8). The node *ν* is referred by the leaf *h*_6_, or by its index in the permuation: *σ*[2] = 6. The subtree rooted at the node *ν* is the interval *σ*[0 : 5] of the permutation *σ*.

### 3.2. Recombination and 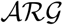 encoding

*A subtree* rooted at a node *ν* in a tree *T* is a subset of *T* which includes all descendant lineages of *ν*. In terms of planar order, it is a semi-open interval [*σ*(*k*), *σ*(*j*)) of the distance vector where *d*_*σ*(*i*)_ (*i*: *σ*(*k*) < *σ*(*i*) < *σ*(*j*)) is less or equal than some given value *D* and such that

- *k* = 0 or *d*_*σ*(*k*)_ > *D*.
- *j* = *M* or *d*_*σ*(*j*)_ > *D*.

A recombination corresponds to a prune-and-regraft operation: one allows to cut an edge and then to glue the cut subtree anywhere on the remaining tree above the cut point.

To keep the planar order, we need to copy an entire block of values of permutation *σ* and of distance vector *d*_*σ*(*i*)_ corresponding to the subtree affected by the recombination (see Figure 2 for the example). To encode this operation we need only three values:

- index of the internal node corresponding to the cut subtree;
- index of the insertion position;
- regraft height: the new value of *d_σ_* between the insertion position and the following row.

**Figure 2.**
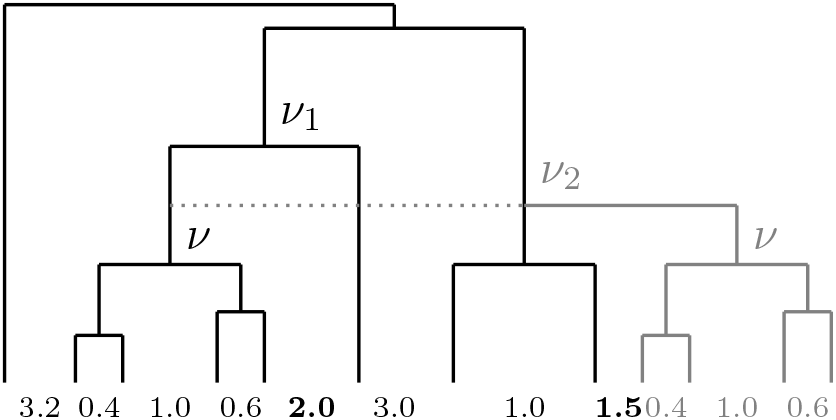
Moving a subtree *ν*: the node *ν*_1_ is destroyed by deleting the corresponding distance value 2.0 (highlighted with bold font), the whole block of distances (0.4,1.0,0.6) is moved to the new place of the distance vector and the new distance value 1.5 is added before this block to create the node *ν*_2_.

## 4. 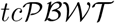 data structure

The core idea of our approach is to use a data structure similar to 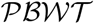 which would naturally carry a lot of structure in itself and would allow fast manipulations with the data and its 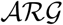. Tree consistent 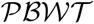, or shortly 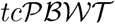, is a modification of the 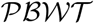 rule. The key difference of 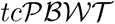 from the original 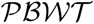 is that rather than moving all the 1’s to the bottom of a column in 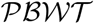 (see section 2.3), in 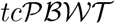 we just require that all the 1’s are contiguous in the new column. A consequence is that in the absence of recombinations it converges to the planar order of haplotypes relatively to the true genealogical tree relating them.

### Definition 2.

Define a map *σ* from the set of binary columns 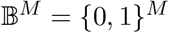 to the set Π of permutations *π* of length *M* to be *tree consistent* if it satisfies the requirements that for any 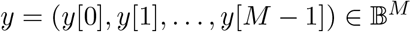

- if for indices *i,j* ∈ [0..*M*), *y*[*i*] = *y*[*j*] and *i* < *j*, then *σ*(*y*)(*i*) < *σ*(*y*)(*j*), that is the order in a subset of zeroes and in a subset of ones is preserved;
- there exist indices 0 ≤ *j,k* < *M* such that *y*_*σ*(*y*)(*i*)_ = 1 if *σ*(*y*)(*i*) ∈ [*j,k*] and *y*_*σ*(*y*)(*i*)_ = 0 otherwise (that is permuted y contains a single block of ones);
- there exists and index *p* ∈ [0..*M*) such that *y_p_* = 1 and *σ*(*y*)(*p*) = *p*.

In other words, every binary column induces a permutation with these particular properties. Evidently, *σ* can be defined as a probabilistic function: for every column it chooses a permutation which satisfies tree consistent conditions, following some probabilistic distribution.

*Example* 1. Let *p* be a minimal index such that *y*[*p*] = 1. We put the block of ones starting from the position of the first appearance of the 1 in the column *y*.

*Example* 2. Let *p* be the beginning of the (first) biggest block of ones in *y*. We keep the position of the biggest block and we move all other ones to it.

Define by induction a sequence of permutations *a_k_* of length *M*, where the permutation that relates *a_k_* to *a*_*k*+1_ is obtained by a map *σ* from the currently permuted column of *H*, *y^k^* = (*h*_*a_k_*(0)_[*k*], *h*_*a_k_*(1)_[*k*], *h*_*a_k_*(2)_[*k*], …, *h*_*a_k_*(*M*−1)_[*k*]). We begin the induction by setting *a*_0_(*i*) = *i*, then *a*_*k*+1_ = *σ*(*y^k^*)*a_k_* where always 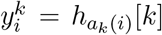. Note that the order *a_k_* at a given site depends on the values of *H* at previous sites, and that we can define a final order *a_N_* after applying maps for all columns. We call the sequence of permuted columns *y^k^* a 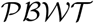. The relation to the standard Burrows-Wheeler Transform [1] for a single sequence is described in Durbin (2014) [2]. If the map *σ* is tree consistent, then we call this transformation a tree consistent 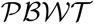.

We call this type of map and 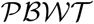 tree consistent because if the haplotypes are related by a simple tree then the ordering converges. Assume that the haplotypes *H* derive from a region without recombination, so that they are related by the same tree at all sites. The values in a column at a site correspond to a mutation on some edge of the tree, and the set of indices taking value 1 at that site will be all the leaves under this edge. If the haplotypes are planar ordered, then each site will have a single block of ones. Tree consistent map will not change the planar order.

We now consider what happens if the *H* derives from a tree, but the initial order of haplotypes is not planar ordered.

### Lemma 1.

*Let H be a set of haplotypes derived from mutations on some coalescent tree without recombination. Consider a* 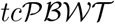 *derived from H. Then for any* 0 ≤ *i* < *k, the column i of H permuted by a_k_, that is (h_a_k_(0)_[i], h_a_k_(2)_[i],…, h_a_k_(M−1)_[i]), contains only one block of ones*.

This follows because each permutation *σ*(*y*^*k*+1^) will not break the block of ones established by *σ*(*y^k^*), although it can either move this block or permute some haplotypes within it. This lemma remains true only for coalescence without recombination. We will discuss what we can say about recombinations later. A direct consequence of Lemma 1 is

#### Proposition 1.

*If a mutation occurred on each edge of the tree, then the final order a_N_ of a* 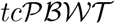 *will constitute a planar ordering of the tree*.

Tree consistent 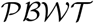 therefore provides a linear time solution to the perfect phylogeny problem [4], page 35. In the presence of recombinations, planar order is destroyed but local trees are still highly correlated. Recombinations will act by splitting blocks of ones. From information theory point of view, in our problem mutations are the source of information and recombinations are the source of chaos. In our algorithm we use a greedy approach for tree transformations. Suppose that we have a local phylogenetic tree *T*_*k*-1_ at a site *k* − 1 and the values of alleles *H_k_* at site *k*. We split *T*_*k*-1_ into maximal subtrees such that values of H_k_ are constant within every subtree. We call this *tree reduction*. For every non-binary node *ν* we create a new child node *μ*. Then all the subtrees rooted at *ν* and carrying derived allele are transferred to the node *μ*. This results in a new tree 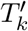 (*tree refinement*). If *H_k_* is still not consistent with 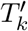, we create SPRs (subtree prune-and-regraft operations) to transform 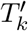 into a tree *T_k_* so that it is consistent with *H_k_*. We consider different strategies to create those SPRs.

There is one more difference of our inference compared to the common approach to 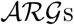. 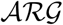 topology is a directed acyclic graph. Our method does not guarantee that the resulting graph does not have cycles. There are two reasons for that. The first reason is that it is faster, though we know a modification of our algorithm with conjectured complexity (*NM* log *M*) which creates graph without cycles. The second reason for allowing cycles is that such approach allows to “forget” errors faster while the algorithm moves along the genome.

## 5. Scalability

To demonstrate the scalability of the algorithm, we use the implementation by V. Shchur based on the planar order data structure. We simulated 100, 1000 and 10000 and 100000 haplotypes with 100000 SNPs using the coalescent simulator *scrm* [12]. The results of the runtime of the algorithm is shown in the table 1 and on the plot 3. The runtime indeed grows linearly in the number of haplotypes.

**Table 1.** Running time in sec of 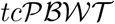 with sets of 100,1000,10000 and 100000 and 10^5^ segregating sites compiled with gcc -O3 option and computed on Intel Xeon CPU E5-2620 v2 2.1 GHz

| Number of haplotypes | $10^2$ | $10^3$ | $10^4$ | $10^5$ |
| --- | --- | --- | --- | --- |
| User time | 0.392 | 2.372 | 21.37 | 251.901 |
| System time | 0.003 | 0.026 | 0.238 | 2.735 |

## 6. Estimating accuracy of topologies

We simulated 100 haplotypes with 50000 SNPs using *scrm*. We estimated trees at each SNP using implementation of our method by by N. Valimaki and V. Shchur. We used different metrics to compare ground-truth and inferred local trees. The main problem with the verification is the computation complexity of the problem of comparing a pair of trees. We need to compare many pairs of trees to evaluate the quality of inference. We chose quartet distance which is one of the classic measures for comparing phylogeny trees. We also developed our own methods for estimating inference accuracy: cluster sizes induced by pairs of leaves.

### 6.1. Quartet distance

We compared inferred trees with trees generated by *scrm* by computing quartet distance between each pair of corresponding trees. Quartet distance is a distance between two (phylogenetic) tree topologies with the same set of leaves. It is defined as the fraction of quartets (sets of 4 leaves) which are related by different subtrees in the two trees under consideration.

We used software *qdist* [7] to compute quartet distance between ground-truth and inferred trees at 50000 sites and mutation to recombination rate ratio equal to 1. The mean quartet distance between these trees is 0.23 with standard deviation 0.09. We present the behaviour of the quartet distance along a fragment of simulated haplotypes at figure 4.

### 6.2. Cluster distance

Cluster distance C for two haplotypes *h_i_*, *h_j_* (*i* ≠ *j*) is the number of leaves under the most recent common ancestor of *h_i_* and *h_j_*. In the 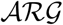 cluster distance between two haplotypes is a function of site 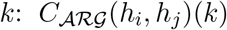. Notice that the set of cluster distances between a given haplotype h_k_ and all other haplotypes *h_i_*(*i* ≠ *k*) define the tree topology completely.

From simulations we know the true underlying 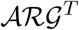 (in the form of the set of local trees). The behaviour of 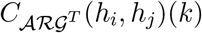 and 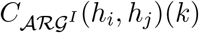 on a fragment of chromosome is plotted in figure 5. Notice that our method captures principal changes in the cluster sizes. To compare these functions numerically, we chose two approached. Firstly, we computed the correlation of functions 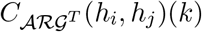 and 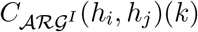 (see table 2). We made these computations with 100 haplotypes and recombination to mutation rates ratios of 1 : 1, 10 : 1 and 100 : 1. Secondly, we compute the the distribution of points with coordinates 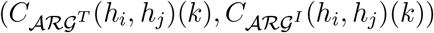 and compute the linear regression of this function (table 2). We plot these distributions for mutation rates ratios of 1 : 1 (figure 6) and 10 : 1 (figure 7). The first distribution shows strong correlation between inferred and true cluster sizes.

**Figure 3.**
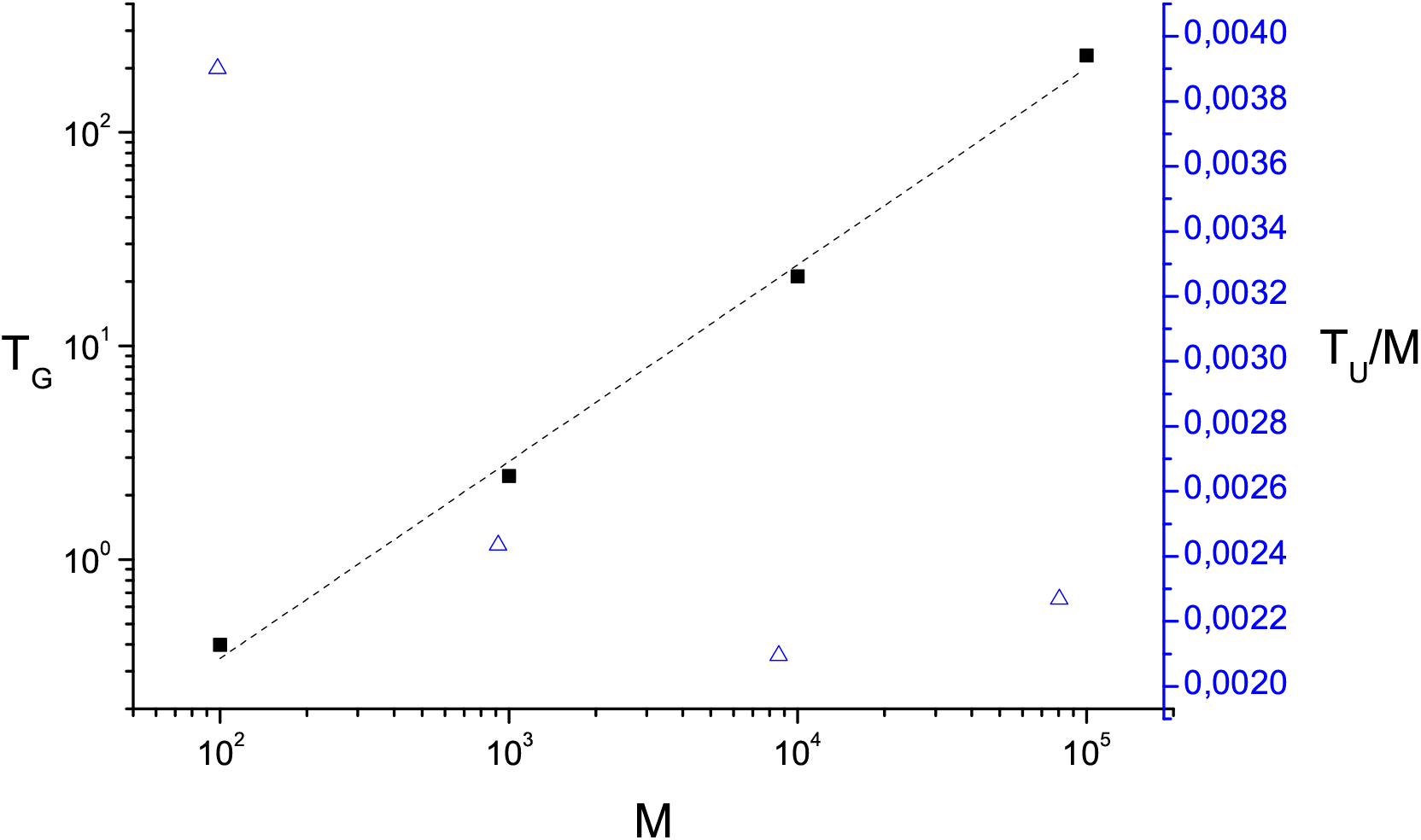
User time in sec of 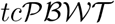 for M = 100,1000,10000 and 100000 haplotypes (solid black squares, left Y-axis). User time normalised to the number of haplotypes M (blue triangles, right Y-axis). Implementation can be found at repository https://github.com/vlshchur/argentum. Computed on Intel Xeon CPU E5-2620 v2 2.1 GHz.

**Figure 4:**
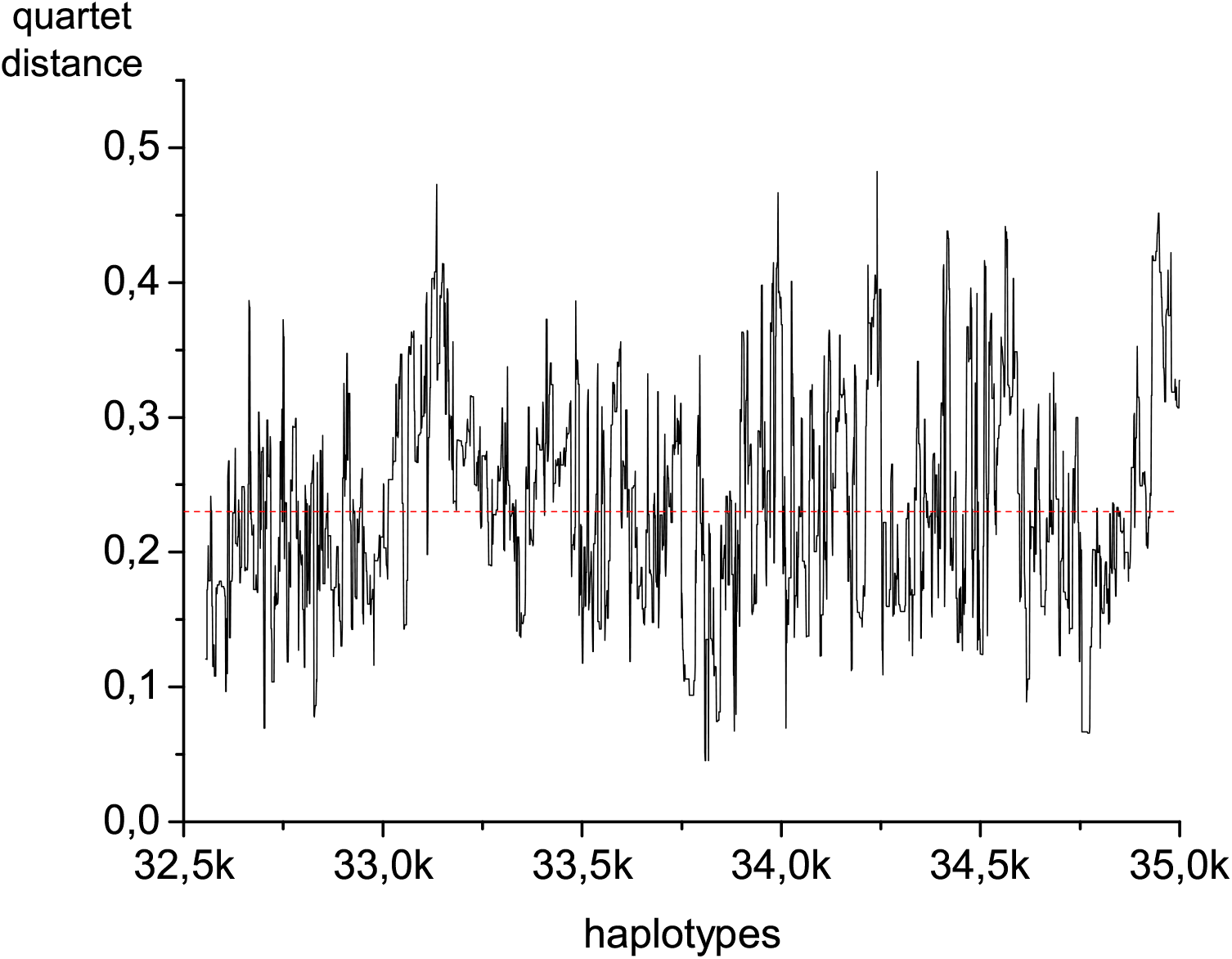
Behaviour of quartet distance between *scrm* and 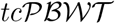 trees along a fragment of haplotypes with 2500 SNPs.

**Figure 5.**
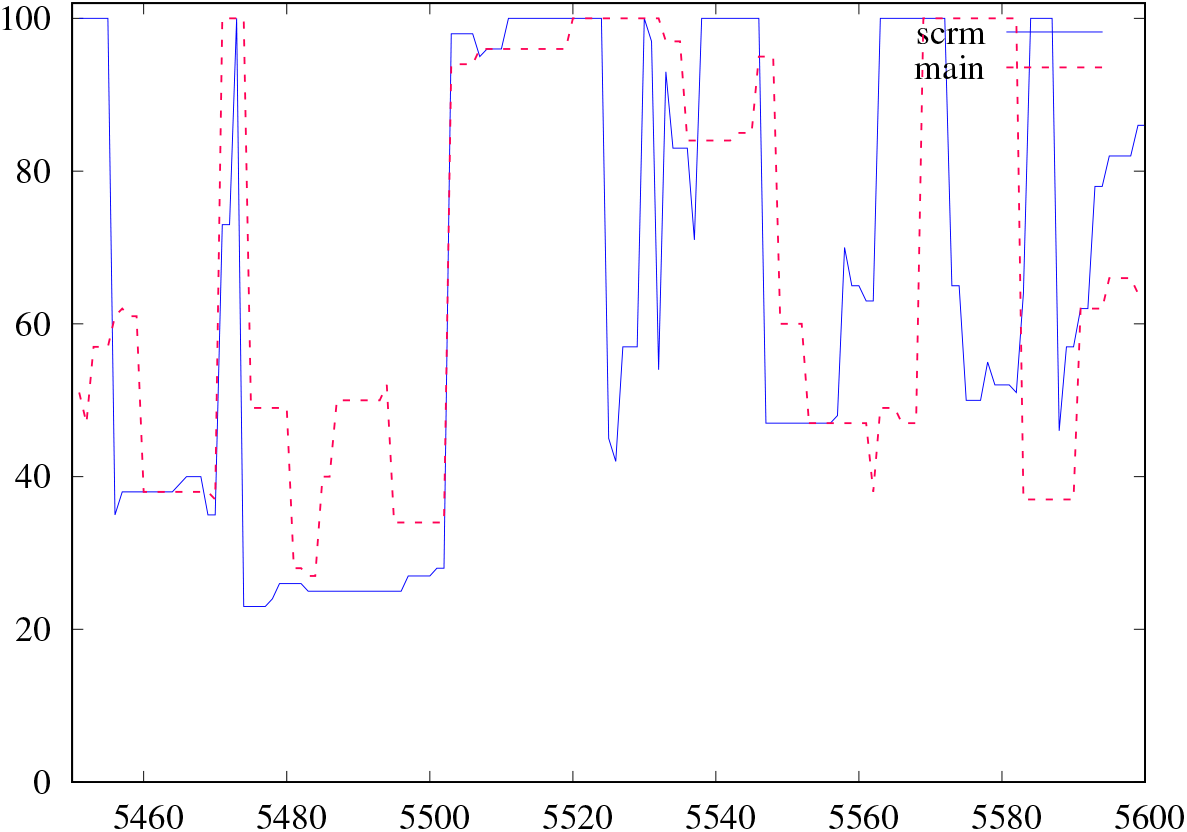
Cluster distance as a function of site for ground-truth 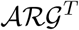 (scrm) and inferred (main) 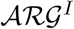. Mutation to recombination rate ratio is 1.

**Figure 6.**
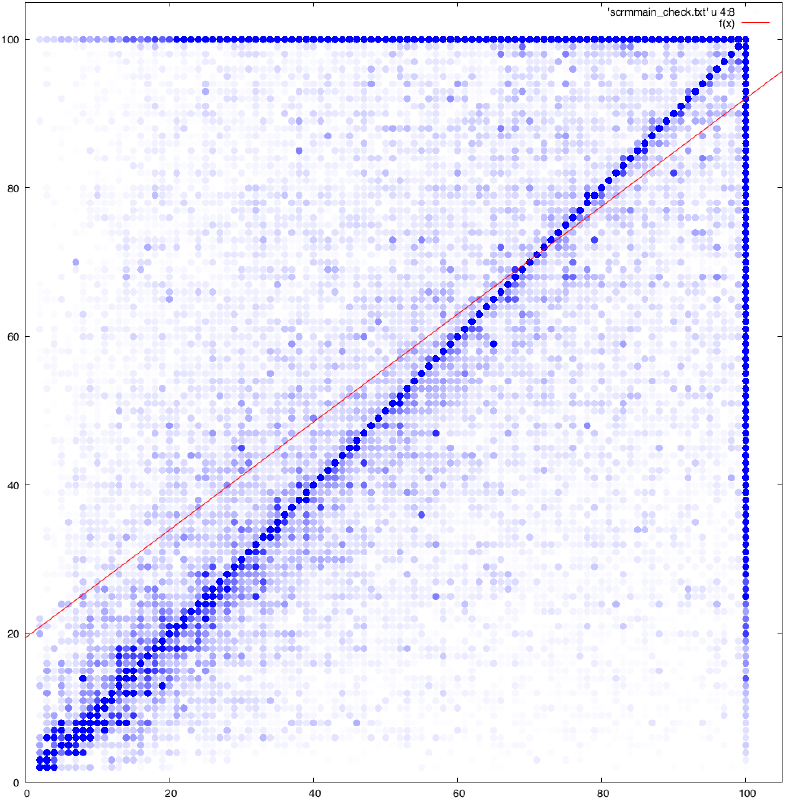
Simulated cluster size against inferred cluster size. Linear regression has slope 0.73. Recombination to mutation rates ratio 1:1, 100 pairs

**Figure 7.**
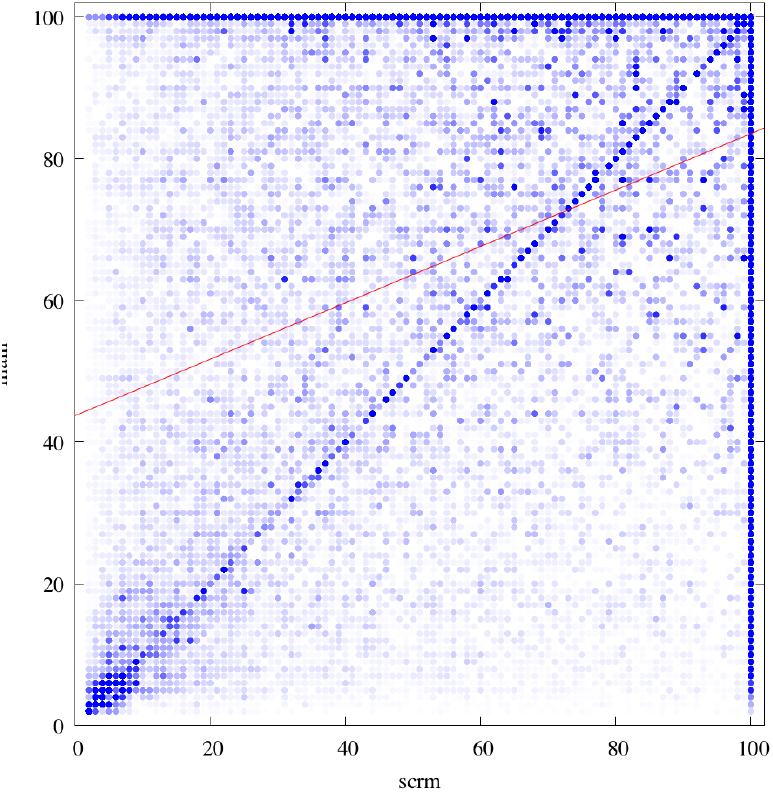
Simulated cluster size against inferred cluster size. Linear regression has slope 0.39. Recombination to mutation rates ratio 10 : 1, 100 pairs

**Table 2.** Mean correlation of cluster size function for true and inferred 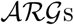 for over all pairs of haplotypes. Linear regression slope for the distribution of cluster sizes in 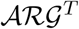 against 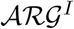.

| Recombination to mutation rate | 1 : 1 | 10 : 1 | 100 : 1 |
| --- | --- | --- | --- |
| Correlation of $C_{\mathcal{ARG}^T}$ and $C_{\mathcal{ARG}^I}$ | 0.7437(2) | 0.4161(2) | 0.2048(2) |
| Linear regression slope | 0.7306 | 0.3877 | 0.1783 |

## 7. Exploratory analysis based on 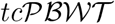

In this section we discuss two approached for exploratory analysis of the data based on the local tree topologies. We applied 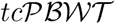 to the humans from 1000 Genome Project (Phase 3). We also analysed simulated data.

### 7.1 Demographic inference from 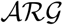 cluster density

#### Definition 3.

*Cluster density D_clust_* of two haplotypes is a vector < *υ*_2_, *υ*_3_,…,*υ_M_* > of length *M* – 1 where *υ_i_* is the number of sites where local cluster size equals to *i*.

Obviously, for more closely related samples the density is higher for smaller values of cluster sizes.

We managed to find the expected distribution of cluster density for a neutral population. Consider the average (over the whole population) cluster density 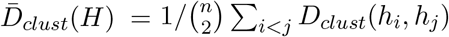.

#### Theorem 1.

^1^ *If the haplotypes of the set H are fully exchangeable, then the expectation* 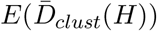 is

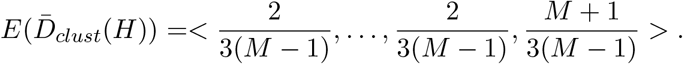

We applied our algorithms to the 1000 Genome Project data (5008 haplotypes). We computed cluster density for all pairs of individuals in the set. Here we present the comparison of British (GBR) to Finnish (FIN), Tuscany (TSI), Chinese (CHS) and Yoruba (YRI) populations, where in each case we take average over all pairs (see figure 7.1). As expected, Yoruba is the most distant population from British. European populations tend to be much more similar than Asian and African. British population demonstrates more similarity to Finnish population than to Italian population based on cluster sizes from 5 to 30. Some lack of similarity between British and Finnish populations compared to Italian population might be explained by the recent bottleneck in Finns, hence they tend to cluster within their own population more frequently. Notice that in the analysis we should take into account the total number of representatives of each population, because it might affect the final distribution.

### 7.2 Demographic inference from 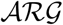: imbalance density

Cluster density allows to compare two particular haplotypes. We also introduce another summary statistic which is called *imbalance density*. Imbalance density is designed to compare two populations. Let the set of haplotypes be divided in two populations *P*_1_ and *P*_2_: 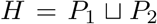. For each branch *B* of each local tree compute the quantity

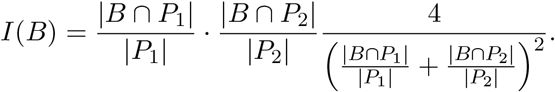

#### Definition 4.

*Imbalance density* is the vector 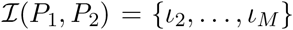 such that *ι_k_* is a mean value of *I*(*B*) over all branches of all local trees possessing *k* leaves:

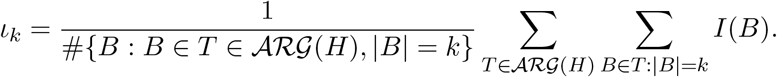

We simulated three sets of haplotypes with different demographic histories. Each set contains two populations with 50 haplotypes in each. The first two scenarios had a split 250 and 500 generations ago respectively (with the normalising effective population size 10000). In the third scenario there was a constant migration between the two populations, but their never merged into a single population.

We plot imbalance density for those scenarios at figure 9. The solid lines are distributions based on simulated trees. The dashed lines are distributions inferred by 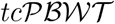. Qualitative analysis of shapes of imbalance density shows that this statistics can distinguish the three demographic scenarios described above. 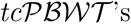 prediction is rather close to the true answer.

**Figure 8:**
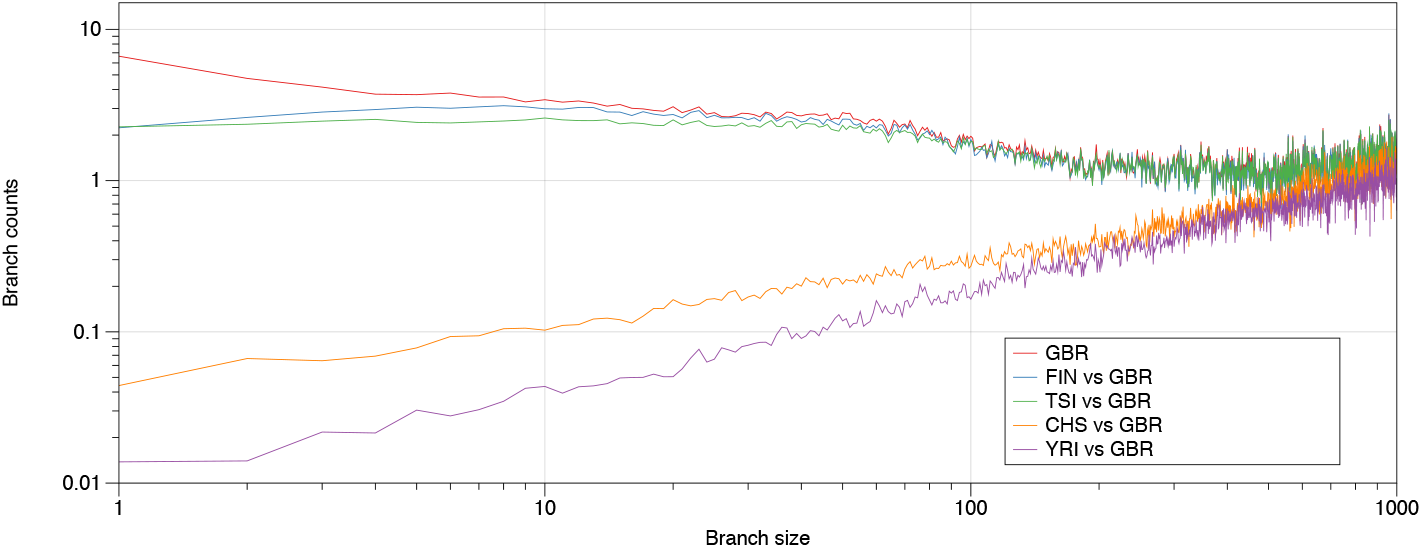
Cluster density for British (GBR) population compared to themselves, Finnish (FIN), Tuscany (TSI), Chinese (CHS) and Yoruba (YRI) populations

**Figure 9.**
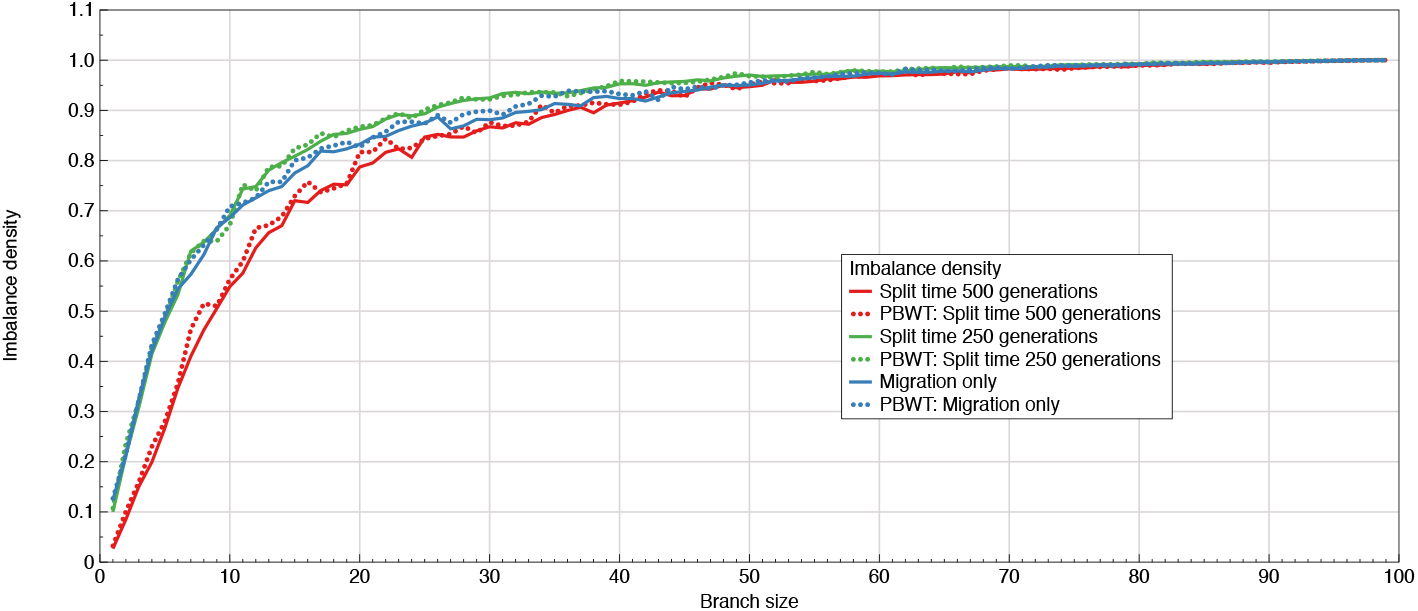
Imbalance density for simulated data.

## 8. Probabilistic model

In this section we will suggest a probabilistic model over possible 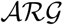 topologies and we will discuss time estimation for 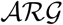 nodes. In our approach, we consider only coalescence nodes of an 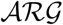 and we do not add recombination nodes in the resulting graph.

Given a set of local trees related by SPRs, one can convert them into a single graph. The nodes and edges shared between adjacent trees (in other words those, which are not destroyed by SPRs) are identified. 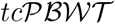 generates such a set of trees and SPRs, so it can be easily converted into such a graph. We will refer to such a graph as 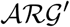 (see figure 10). The difference with the regular 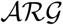 is that we have only coalescence nodes, but we exclude recombination nodes. This data structure is closely related to coalescence records format [6].

**Figure 10.**
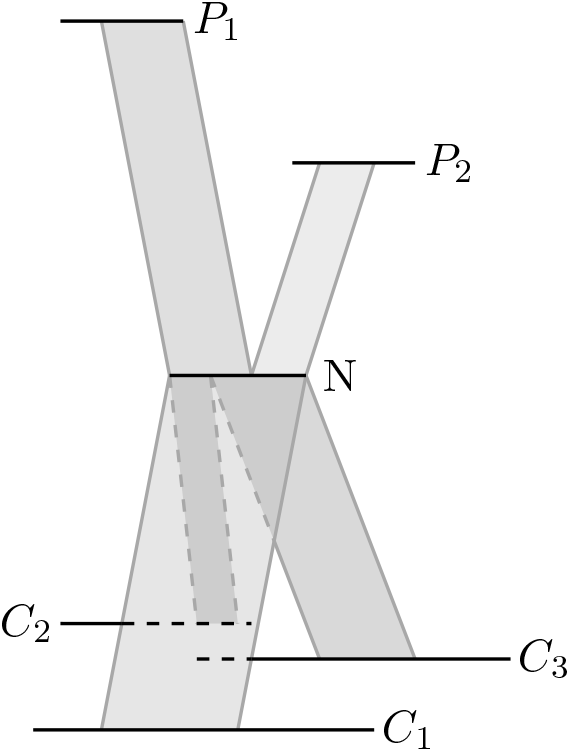
Example of a node in the 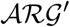: the node *N* has two parents *P*_1_ and *P*_2_ and three child *C*_1_, *C*_2_ and *C*_3_.

Then we extend the standard model for mutation and recombination event on trees to the whole graph. Let *G* = (*V, E*) be an 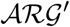. Let an edge *e* of *G* span a certain genomic interval *I*(*e*) and a certain time *T*(*e*). The number of events *N_e_* on the edge e is distributed accordingly to Poisson distribution with the rate *λ_e_* = *μ* · *I*(*e*) · *T*(*e*) where *μ* is the mutation rate.

Hence the likelihood of the data given topology of the graph G with mutations and recombinations assigned to edges is

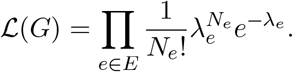

Hence the maximum likelihood problem can be translated to the problem of mathematical programming in the following way. We pass to the log-likelihood

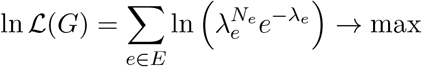

or

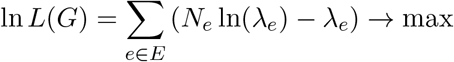

Notice that the function *f*(*λ*) = *N* ln(*λ*) – *λ* is unimodal with the maximum at *λ* = *N*.

After some transformations we end up with the following problem of mathematical programming

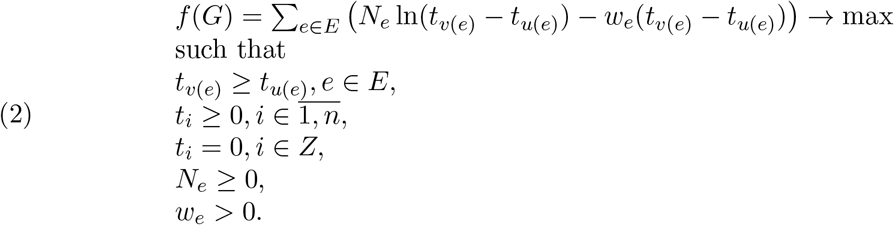

Unfortunately, we did not find solution of this problem. Maybe, 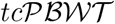 heuristic approach need some modifications or extension of this likelihood function, so further study is needed.

It would be also desirable to introduce the concept of local likelihood function of a tree. The idea comes from the following observation: while inferring tree topologies, at some fragments we get reasonably good estimates, in others there are many errors. Hence, the full likelihood of an 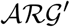 is not a good measure of the inference quality of a tree at a particular site. Instead it would be interesting to consider local tree with “extended” edges: the edges from the 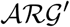 and including all the mutations from all nearby sites belonging to those extended edges. This would allow us to integrate over local trees at each site independently.

## 9. Discussion

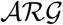 is arguably the most interesting data structure in evolutionary biology. If we were able to estimate it precisely enough, we would get access to the full information about evolutionary forces shaping populations: e.g. changes of effective population size, split times, migration rates, selection coefficients. However, inference of 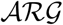 is a very complex problem because of the complex state space and many unobserved parameters, especially recombinations.

We have suggested a partial solution of this problem, which allows us to relate chromosomes by highly correlated tree topologies. Our method scales linearly both in the number and in the length of sequence. We show that these tree topologies can capture both global population structure and local tree structure. For the problem of global population structure inference we introduced two new statistics defined on sets of local trees – cluster density and imbalance density.

We also discussed potential ways of extending our approach to enable probabilistic analysis and estimation of node times in the graph. Currently this has more theoretical and conceptual than practical value, but we hope to develop these ideas in future.

## 10. Acknowledgement

We thank Stephen Schiffels for fruitful discussions and the idea of imbalance density. We thank Niko Välimaki for working on the implementation of 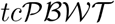 and exploring different subalgorithms. Liliia Ziganurova was supported by grant RSCF 14-21-00158. Vladimir Shchur and Richard Durbin were supported by Wellcome grant WT098051.

## Supplementary materials

### 1. 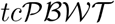 framework

Our proposed 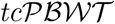 approach for 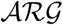 inference consists of the following steps.

- At the first stage it generates local tree topologies from the data.

- Forward run. The algorithm traverses the genome position by position. At each position it updates the local tree by identifying the minimal number of subtree prune-and-regraft (SPR) operations needed to explain the data. The number of SPRs is minimised relative to a single site following certain rules and restrictions. Different modifications of those rules are suggested. In particular, the process can be randomised for sampling purposes and statistical inference.
- Backward run. The algorithm is not symmetric in terms of the direction of traversing genomes. Backward run is used to reduce this effect. For a given site, the local tree is built based only on the information from one side of the genomes (based on prefixes) during the forward run. The backward run allows to refine the local trees by using suffixes too. As a result, nodes inferred from the opposite side of genomes are added to local trees.
- A modification of the algorithm exists which allows to run 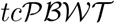 from left to right and from right to left and then “merge” the inferred sets of the trees.
- The set of local tree topologies is converted into a single graph by identifying the shared tree nodes.
- We present a tractable probabilistic model for the resulting graph, which allows to estimate times of its nodes.
- We also suggest a concept of local likelihood of the data. One can generate a large set of 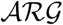 topologies. At each genomic position one gets a sample of local trees (induced by those 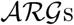) with their likelihoods. Then one could integrate over this sample for their inference.

We succeeded to show the performance (both scalability and reasonable accuracy) of the first step which generates local tree topologies. We establish a probabilisitic model which is consistent with SMC [9] model, and state a corresponding optimisation problem. Unfortunately we do not have a fast algorithm which could solve it yet.

### 2. Data compression results for 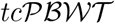

We performed a comparison of a compression performance of 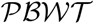 and 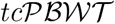 on a simulated set with 1000 haplotypes and 148120 segregating sites (table 2).

We note that the original 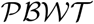 introduced by R. Durbin is not tree consistent. Its map *σ*, which takes the entire block of ones to the end of the column, satisfies the first two requirements given above, but not the third. When we apply the original 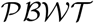 to a tree ordered set of haplotypes then the permutation typically changes at each site even for a simple single tree genealogy. That is why we can expect an improvement in the compression rate by 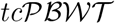 compared to 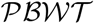.

**Table S1.** Comparison of 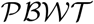 and 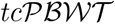 compression performance. The columns “blocks” show the total number of blocks of ones which appear in a column k + 1 relatively to 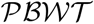 or 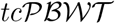 built at position k. File sizes are shown in bytes. The number of blocks in 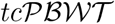 is less by 21% for simulated data.

| Method | Blocks | File size |
| --- | --- | --- |
| $\mathcal{PBWT}$ | 370302 | 1,207,976 |
| $tc\mathcal{PBWT}$ | 306126 | 1,074,540 |

### 3. Combinatorial 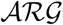 inference

In this section we will discuss some core algorithms of our approach for 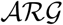 generation. Those algorithms are based on 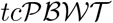 data structure and planar ordering format. The 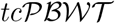 can be considered as a certain number of basic rules for data transformation and tree refinement. Planar ordering is a format which allows fast operations and good optimisation of software implementations as well as compact data representation.

We call a leaf with an allele *υ* (*υ* = 0,1) shortly a *ν-leaf*. A *υ*-subtree (*υ* = 0,1) is a subtree such that all its leaves are *υ*-leaves. For a node *ν* we denote the set of all *υ*-leaves rooted into *ν* by *L_υ_* (*ν*).

We often refer to a node and a corresponding subtree interchangeably.

#### 3.1 Tree reduction

The local trees *T*_*k*-1_ and *T_k_* are highly correlated, hence *T_k-1_* induces a lot of structure at the site *H_k_*. That means that the haplotypes carrying the mutation at the site appear only in a few subtrees of the local tree at the preceding position. We use this property to compress the tree *T*_*k*-1_ relatively to the binary column *H_k_* and call this process a *tree reduction* (see Figure 3.1 TODO). This algorithm is linear in M and allows us to perform further computations in a smaller space. In fact, this idea is close to the one which enables an efficient 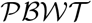 compression (see [2]).

A *maximal constant subtree ν* of a tree *T*_*k*-1_ relatively to the site *H_k_* is a subtree such that all haplotypes in it have the same allele at site *H_k_*, and any subtree 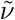 such that *ν* is its proper subset 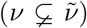 includes haplotypes with different alleles. Maximal constant subtrees induce a disjoint subdivision on the set of leaves of *T*_*k*-1_ uniquely.

A *reduced tree* 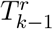 is obtained from the tree *T*_*k*-1_ and the site *H_k_* by substituting all the maximal constant subtrees by one leaf with corresponding allele. The distances for 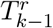 are naturally induced by distances in the full tree *T*_*k*-1_.

The algorithm is written in details in Algorithm 1. The planar ordering of *T*_*k*-1_ guarantees that every subtree appears as a single interval. Relying on this property, the tree reduction algorithm identifies stretches with constant allele values. Each such stretch is then represented as a maximal set of subtrees through the function `Parselnterval()`.

**Figure S1.**
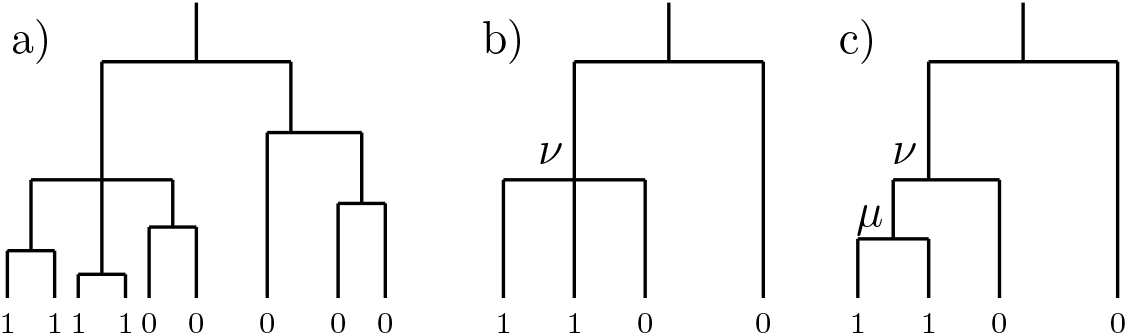
Tree reduction and tree refinement. a) An example of a tree *T*_*k*-1_ with a site alleles *H_k_*. b) A reduced tree 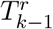. c) A refined tree with a new node *μ* under the node *ν*. Now *μ* is a parent node of 1-subtrees previously rooted at *ν*.

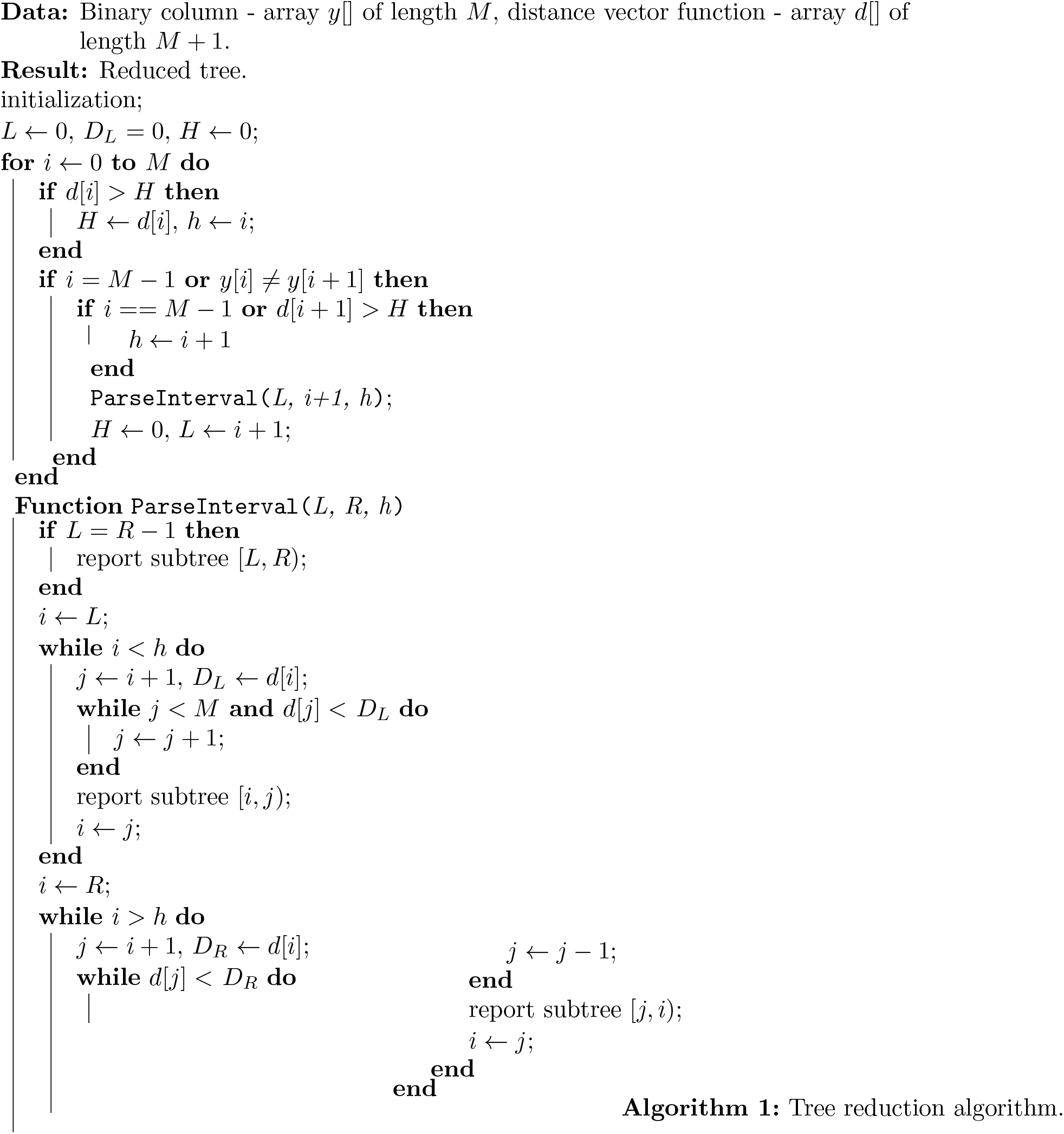

**Figure S2.**
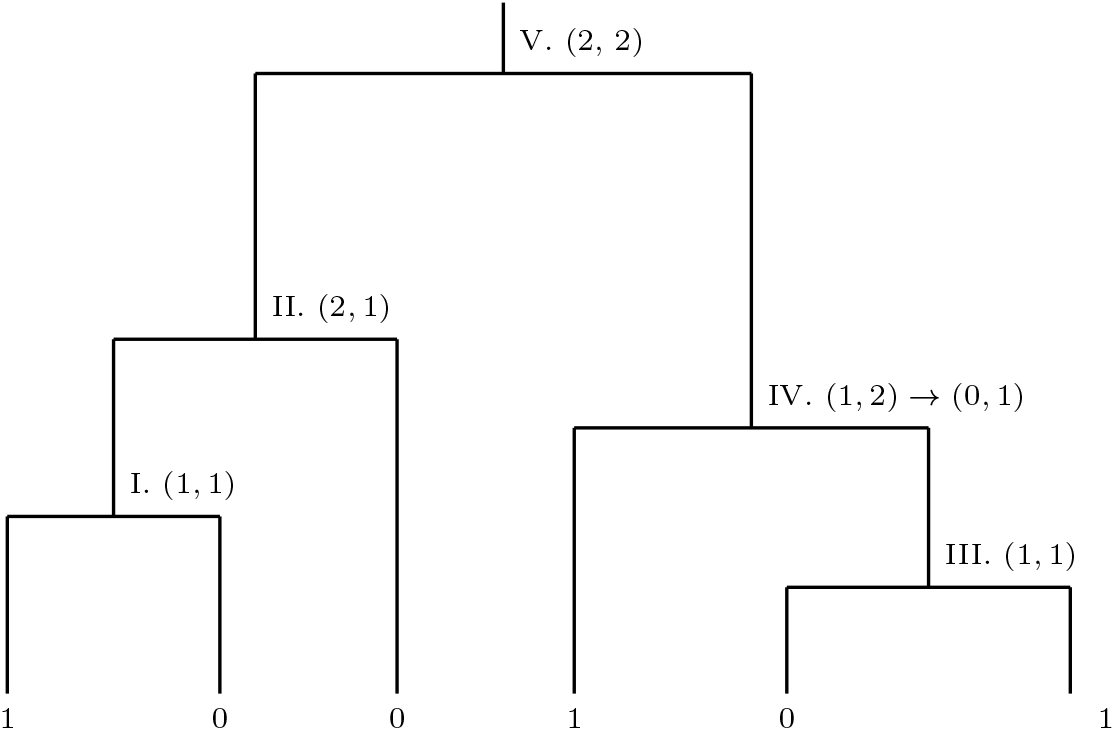
Reducing the number of recombinations. Corresponding stack for node V.: Push V, go to node IV. Push IV, go to node III. Resolve III. Pop from stack node IV. Resolve IV. Pop from stack node V.

#### 3.2 Tree refinement

At this stage in our approach we try to increase the number of internal nodes of the tree using tree refinement procedure: at every non-binary node we merge together 1-subtrees. This algorithm works on a reduced tree instead of a complete tree (see Figure 3.1 transition from *b* to *c*).

In a reduced tree the algorithm visits all non-binary nodes. If for a node *ν*, the set of 1-leaves *L*_1_(*ν*) contains at least two elements, then a new node 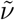 is created. The parental node of 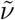 is set to be *ν* and the elements of *L*_1_(*ν*) are reassigned to 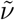 as child leaves.

After this procedure *ν* is still well defined. It has at least two child nodes one of which is 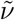. The existence of another leaf follows from the definition of a reduced tree: if all child nodes of *ν* were 1-leaves, *ν* would be reduced to a single leaf itself.

#### 3.3 Reducing the number of recombinations – Part 1

If the tree *T*_*k*−1_ is not consistent with the site *H_k_*, we need to apply SPRs (which represent recombinations) to rebuild the tree. Our procedure uses simple operations on trees in order to reduce the number of recombinations needed to explain the data. At each site it finds the minimal number of SPRs needed to rebuild the local tree to make it consistent with the column. This process does not necessary lead to a global minimum of recombinations needed to explain the data.

For the clarity of exposition we will present the procedure in few steps with introducing more details and elements step by step. Our first goal is to introduce Algorithm 2 which would minimise the number of prune-and-regraft operations on a tree *T*_*k*−1_ to make it consistent with a binary column *H_k_*. Evidently the solution can be found on the reduced tree: to minimise the number of SPR operations one would want to cut maximal subtrees of the tree *T*_*k*−1_. We call this algorithm *MinCut*.

For this purpose the algorithm visits all internal nodes starting from the bottom of the tree and going upwards to the root. It counts the numbers of 0- and 1-leaves below each node. As soon as for a node *ν* the corresponding subtree is incompatible and the number of, say, 1-leaves is larger than the number of 0-leaves, it cuts corresponding 0-leaves and reduces *ν* to a single 1-leaf.

The planar order data structure supports the following linear time realisation of this algorithm. Suppose that we are at position *k* associated with a node *ν*. Let *ν*_1_ and *ν*_2_ be its two child nodes, which are encoded by positions *k*_1_ and *k*_2_ of distance vector and such that *k*_1_ < *k* < *k*_2_. We scan the tree representation from left to right, so by arriving at *k* we already resolve the subtree corresponding to *ν*_1_, but did not enter in the subtree corresponding to *ν*_2_, hence cannot conclude about *ν* too. Hence we push *ν* in a stack and begging resolving *ν*_2_ in the same manner. As soon as *ν*_2_ is resolved, we push *ν* from the stack.

For simplicity we present the algorithm for binary trees. The extension for non-binary trees is rather straightforward.

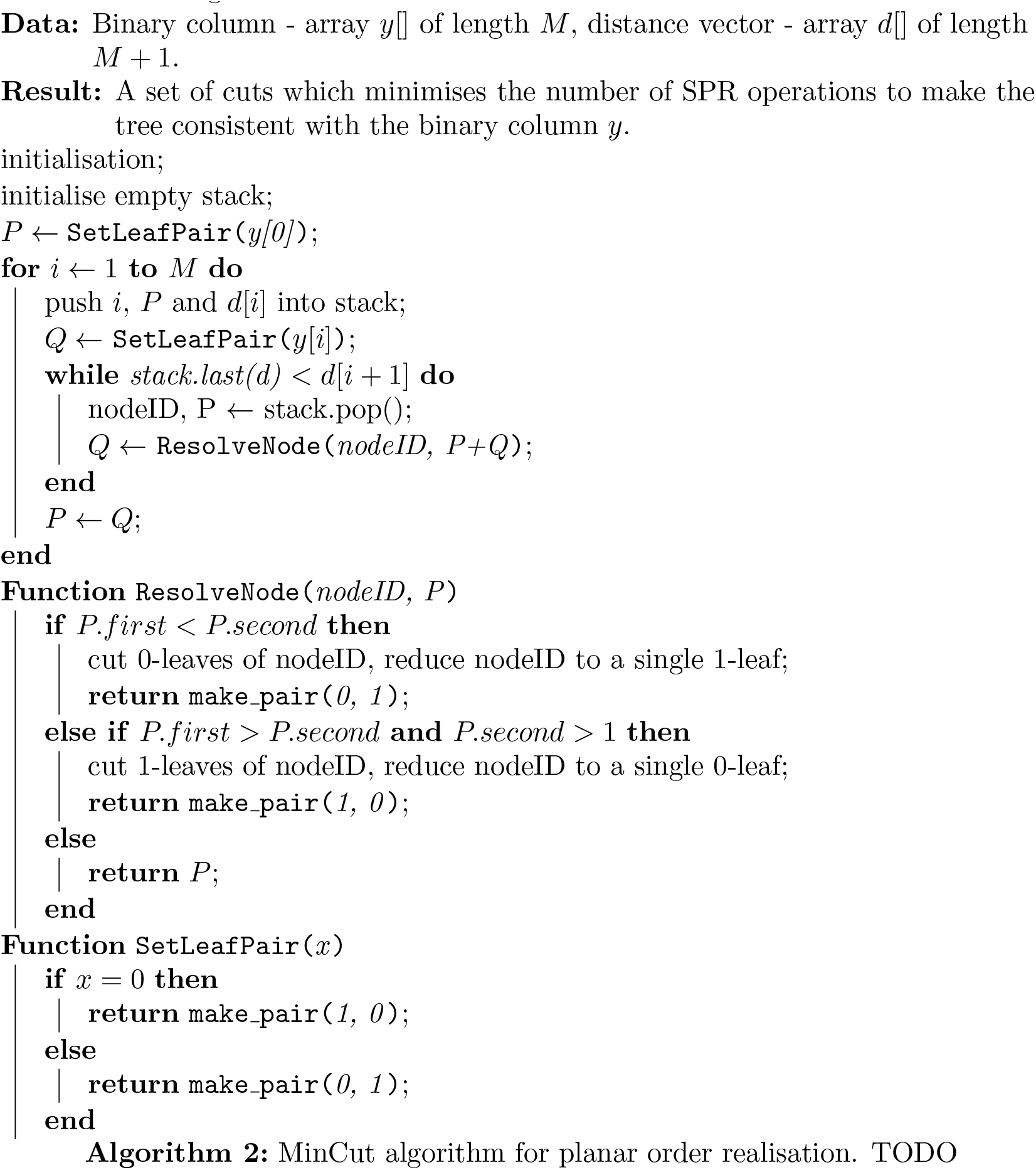

#### 3.4 Choosing new parental node for regraft

If we prune 0-subtrees, we regraft them straight to the root of the tree. Though it might look as a considerable loss in the information about the shape of a tree, we will explain in the next subsection how we struggle against this problem.

For 1-subtrees, we select one *stable subtree*. This selection can be done either deterministically (e.g. to choose a subtree with the maximal number of leaves) or randomly (e.g. with weights proportional to the number of leaves). We will discuss the strategies later on.

All other 1-subtrees are rooted directly to the root of the stable subtree. Such a choice follows from the following partially heuristic (?) idea. On the one hand SPR operation can take a subtree anywhere inside the stable subtree. So regrafting the displaced subtree directly into the root node of the stable subtree allows to relax relations imposed by this node previously. On the other hand, when we apply the SPR operation to a subtree, we keep it as a single object and we preserve its entire structure, that is why we do not destroy root nodes of subtrees affected by recombinations.

#### 3.5 Reducing the number of recombinations – Part 2

Until this moment, we discussed the algorithms which made all the decisions on rebuilding local trees from a single site. Moreover, a local tree *T_k_* used information only about the mutations to the left of the site *k*, though it would be highly desirable to make use of the right side as well. Now we are going introduce the development of the presented data structure to make the decision on the prune-and-regraft operation selection from several columns and to introduce a sort of “looking forward” algorithm without loss in algorithm complexity which remains *O*(*MN*).

We add the following information to our data structure:

- for every SPR operation, a record is created: it specifies the identifier of the displaced subtree, its initial location, its target location in the tree and the locus at which it was applied;
- if a subtree was displaced by a recombination, it gets a tag and a pointer to the corresponding SPR record. A tag is destroyed whenever the parental node of the corresponding subtree becomes binary;
- for every node, one keeps an identifier of the site at which this node appeared in a reduced tree as a 1-subtree.

The basic idea of introducing the tags is the following. When we visit a node *ν* and we see a tagged subtree *ν_t_* rooted at *ν*, we know that there was an evidence from one of preceding columns that the subtree is under *ν* though we do not have enough evidence about the exact placement of *ν_t_* in this particular part of the tree. So as soon as we have more evidences (given, that the new place is still in the subtree under the node *ν*) instead of creating a new SPR operation, we move *ν_t_* to its new position and edit the corresponding SPR record. This process allows to refine the structure of tree without introducing additional recombinations.

For a tagged 1-leaf λ (and a corresponding SPR record *r*_λ_) rooted at *ν* and a node *ν*_0_, which is a descendant of *ν*, we define an operation of *reassignment TODO*. The parental node of λ is changed from *ν* to *ν*_0_ and in the record *r*_λ_ we edit the target from *ν* to *ν*_0_ as well. The reassignment is allowed only if on the path from *ν* to *ν*_0_ all the nodes were created before *r*_λ_(*T*). If a node was created after *r*_λ_(*T*), we need to create a new SPR operation to insert a subtree under this node.

We extend the algorithm from Subsection 3.3. We visit all the nodes of the reduced tree starting from the bottom. Suppose that we already processed all nodes under a non-binary node *ν*. Child nodes of *ν* can be of the following types:

- non-tagged 0-leaf, denote this set by 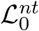;
- non-tagged 1-leaf, denote this set by 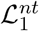;
- tagged 0-leaf, denote this set by 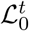;
- tagged 1-leaf, denote this set by 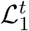;
- there is a draw produced by the algorithm in a non-tagged subtree, 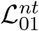;
- there is a draw produced by the algorithm in a tagged subtree, 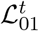;

Both non-tagged and tagged 0-leaves remain on their positions and are not alternated in any way.

Non-tagged 1-leaves are merged in a new node following the procedure described in Subsection 3.2.

A draw in a tagged subtree is resolved by cutting 0-leaves. Then it is considered as a tagged 1-leaf.

Now there are two options.

I. If the set 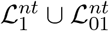 is not empty, we apply the algorithm from Subsection 3.3 to the node ν excluding tagged 1-leaves. If the algorithms reports 1-leaves to be cut, then tagged 1-leaves are reassigned to them randomly.
II. If there set 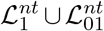 is empty but in the set 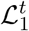 there are at most 2 leaves, we choose one of them randomly to be an analogue of a stable subtree, and then we reassign the rest of the leaves to the selected one.

### 4 Choosing recombination stable branch based on supplementary tree

During the first phase, our method scans the haplotypes in one direction (forward run). We want to make use of the information to the other side of the given site for our inference. Let us firstly generate an 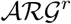 with our algorithm while scanning our data from right to left. Now we modify the left-to-right scan for the inference of 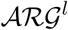. We will use the upper indices *r* (right) and *l* (left) in the notations of this section to highlight by which 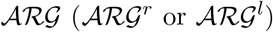 it is induced.

Suppose that a tree 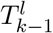 is incompatible with the site *H_k_*, so we need to apply a recombination.

Then we apply a MinCut algorithm (Algorithm 2) to identify which branches to cut. Let 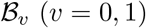 be the set of *υ*–branches to be cut and 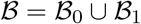.

A 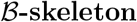-**skeleton** of 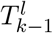 is the tree 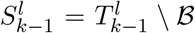, which is the remainder of the tree after removing all branches from 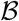. By definition 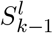 also includes the nodes (which could be of out-degree 1!) which correspond to the nodes of joining with branches from 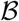.

Now we define a metric on a 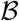-skeleton. The topological length of the path between two nodes in a graph is the number of edges in this path. For each node *ν*, its depth *ρ*(*ν*) is the topological length of the longest path to its descendant (the depth of leaf is 0 hence). Set the length of an edge *e* between two nodes *ν*_1_ and *ν*_2_: |*ρ*(*ν*_1_) – *ρ*(*ν*_2_)|.

For each *h_i_* such that *h_i_*[*k*] = 0 we compute for the right tree

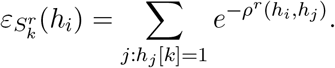

Now choosing each 1-branch from 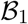 as a stable branch subsequently, we compute the same quantities for 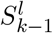 skeleton. The final choice of the stable branch is those which minimises the mean square error

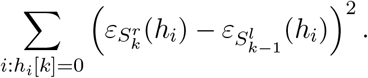

As soon as this choice is done, we add the resulting branch to the skeleton, recompute metric and use this procedure to find the best place for each 0-branch separately.

### 5. Proof of the theorem on the distribution of cluster sizes

Consider a population *H* with *M* individuals *h*_1_, *h*_2_,…, *h_M_*. We suppose that at each time point every pair of lineages has equal probability to coalesce. Each given pair of individuals, e.g. *h*_1_ and *h*_2_, defines and induces a cluster as the set of individuals from H which are descendants of the most recent common ancestor of *h*_1_ and *h*_2_. We are interested in the probability distribution of the induced cluster sizes.

This distribution does not depend on the coalescent times or the effective population size history.

#### Theorem 2.

*Under the coalescent with full exchangeability, the probabilities for induced cluster size*

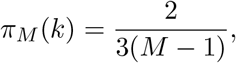

*for* 2 ≤ *k* < *M and*

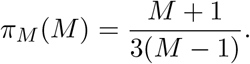

#### Definition 5.

Hierarchical topology of a binary tree is the topology (branching structure) together with the order of coalescences.

It is well-known (see for example J.Wakeley “Coalescent theory” page 82 or Hein “Gene genealogies…”) that the number of different hierarchical topologies of size *M* is

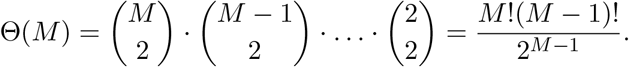

All hierarchical topologies are equiprobable.

### 6. Number of hierarchical topologies with a given subtree

Suppose that *k* leaves appear together in a subtree with known hierarchical topology. How many hierarchical topologies of size *M* contain this subtree “as it is”? We do not allow to change the inner structure of the subtree, in particular we do not allow to add more leaves to it.

#### Lemma 2.

*For a fixed hierarchical topology of size k there are exactly*

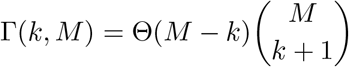

*hierarchical topologies of size M containing it as a subtree*.

*Proof*. Let *n_j_* be the number of coalescences in the reminder of the tree in the time interval when there are exactly *j* lineages in the fixed subtree. Denote 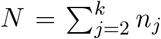. Then the number of trees is

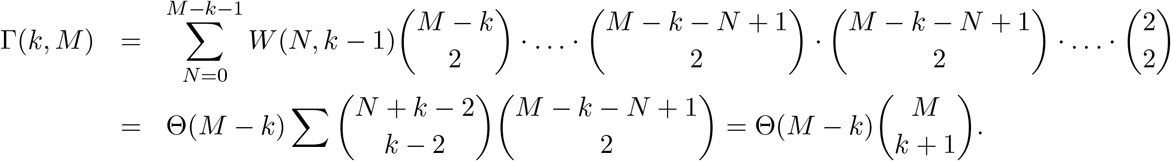

where *W*(*N, k* − 1) is the weak composition, in other words the number of ordered tuples (*n_k_*, …, *n*_2_) such that Σ *n_j_* = *N* and *n_j_* ≥ 0.

### 7. Proof of Theorem 2

#### 7.1. Case of *k* = *M*

In this case at each coalescent event we allow any coalescence except between lineages containing our two selected leaves, which means all except one coalescences. The number of trees where *h*_1_ and *h*_2_ merge only at the root is

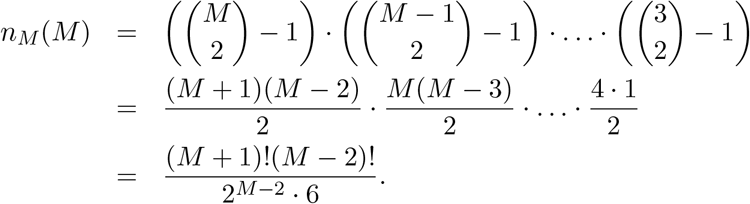

And

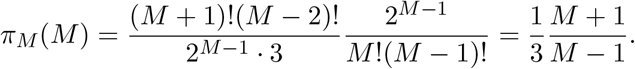

#### 7.2. Case of *k* < *M*

Now we are ready to write and compute the probability that the local cluster density of two given leaves is *k* < *M*. It is given by the following formula

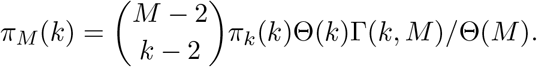

The first binomial coefficient stands for the choice of *k* − 2 leaves which appear in the subtree together with *h*_1_ and *h*_2_. The next two terms count the number of subtrees of size *k* where the lineages of *h*_1_ and *h*_2_ meet at the root. Then we use Lemma 2 to find the number of hierarchical topologies containing a given subtree of size *k*. And finally we normalise by the total number of hierarchical topologies of size *M*.

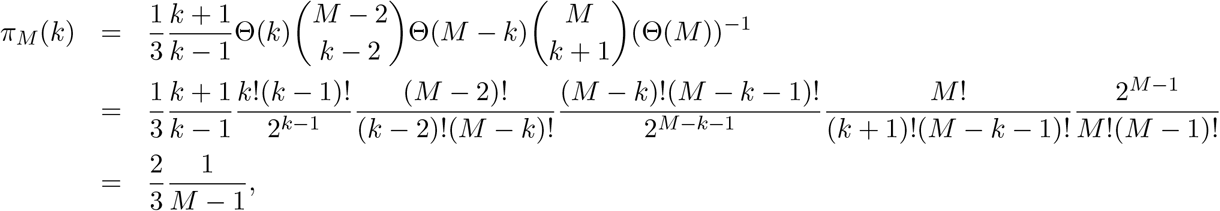

which finishes the proof.

### 8. Probability for two given leaves induce a cluster of size *k*, case of non-hierarchical topology

Now let us suppose that we do not care about the order of coalescences and assume that all topologies (only branching structure, no order on nodes!) are equiprobable.

#### Theorem 3.

*If all leaves are interchangeable, then the probability that the induced cluster size of two given leaves is k equals to*

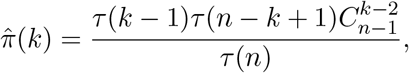

*where τ*(*n*) = (2*n* − 3)!! *is the number of topologies of size n*.

#### 8.1. Number *τ*(*n*) of binary trees with n labeled leaves (see also Hein)

The number of trees with *n* = 2 leaves is *τ*(2) = 1.

Suppose that we have already enumerated all trees with *n* − 1 leaves *h*_1_, *h*_2_,…, *h*_*n*−1_. To obtain all trees of size *n* we add one more leaf *h_n_* to the set of leaves. Now for every tree with *n* − 1 leaves we connect successively *h_n_* to every edge of that tree (including a “free” edge at the root). Connection of *h_n_* to different edges leads to different trees. Hence, *τ*(*n*) = *τ*(*n* − 1) * (2*n* − 3), because a binary tree with *n* − 1 leaves has 2*n* − 3 edges. Hence,

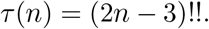

#### 8.2. Number of trees where given leaves are in a subtree with *k* leaves

Let us find the number *σ*(*k*) of trees where two given leaves *h*_1_, *h*_2_ appear in a subtree (not necessarily minimal!) with *k* leaves. We have *k* − 2 free places in such a subtree and we choose them from a pool of *n* − 2, hence there are 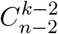 possibilities. Given *k* leaves there are *τ*(*k*) different ways to construct a subtree. Now it remains *n* – *k* leaves and a subtree which we consider as a new leaf. Hence there are *τ*(*n* − *k* + 1) trees which possess a given subtree.

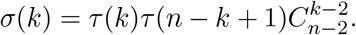

#### 8.3. Probability *π*(*k*) where given leaves are in a minimal subtree with *k* leaves

This is equivalent that two leaves induce a cluster of size *k*. So we obtain

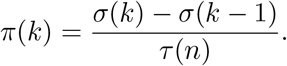

Let us simplify the expression

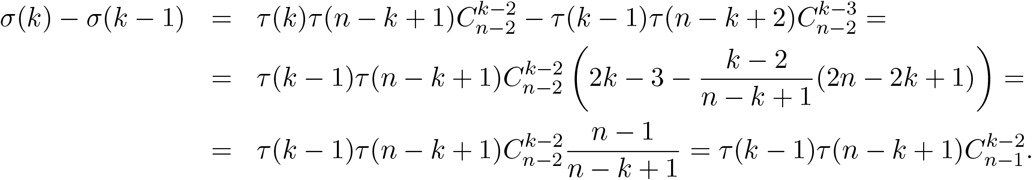

Finally

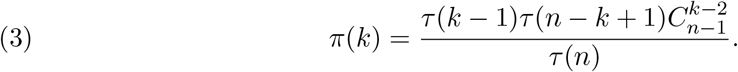

## Footnotes

1 See supplementary materials for the proof of this theorem.

